# Dynamics of electrical resistance of kombucha zoogleal mats

**DOI:** 10.1101/2022.11.03.515122

**Authors:** Andrew Adamatzky

## Abstract

We demonstrated that zoogleal mats of kombucha exhibit spikes of electrical resistance. A kombucha is a sugared tea fermented by a symbiotic community of over twenty species of bacteria and yeasts which produce cellulosic gelatinous zoogleal mats. We recorded electrical resistance of the zoogleal mats via platinum electrodes placed at a distance one centimetre of each other. We found that the mats show temporal variations in electrical resistance in a range 0.13 MOhm to 0.19 MOhm. We discovered spikes of the mats resistance morphologically similar to action potential spikes. Average duration of a resistance spike is 1.8 min, average amplitude is 2.2 kOhm. Average interval between resistance spikes is c. 20 min. The discovered resistive spiking of kombucha mats might indicate on their memfractive properties, and thus open pathways towards prototyping neuromorphic devices with living zoogleal mats.

## 1. Introduction

Electrical resistance of living substrates characterises their physiological and morphological states [1, 2, 3, 4, 5, 6, 7, 8, 9, 10, 11]. By measuring electrical resistance we can determine states of organs [12], estimate roots vigour [13], detect a decaying wood in living trees [14, 15], classify breast tissue [16], assess freeze-thaw injuries of plants [17]. Why did we get interested in electrical resistance of kombucha zoogleal mats?

A kombucha is a tea fermented by a symbiotic community of bacteria and yeasts [18, 19, 20, 18, 21, 22]. The symbiotic community of bacteria and yeasts produces the cellulosic gelatinous mat, also known as biofilm, commensal biomass, tea-fungus, scoby and zooglea. A tea fermented by the symbiotic community allegedly exhibits a range of health beneficial properties [23, 24, 19, 25, 26, 27, 28, 29, 30, 31, 32], however these are outside our current interests. Our interest is in studying kombucha zoogleal mats is threefold.

First, kombucha zoogleal mats can be used as ultra-low cost and fast-to-produce components of future living electronics, sensing and computing devices. We have already demonstrated a potentially unique properties of kombucha mats via analysis of their electrical potential dynamics [33]. We have also shown that acetobacter colonies, which form thick bacterial films, can be used for sensing purposes [34, 35].

Second, kombucha mats are unique symbiotic systems where over sixty species of yeasts and bacteria cooperate [18]. A kombucha is an example of a proto-multicellularity — an organism combined of multiple species each one pursuing a common goal of prolonging a life time of the collective organism. Electrical properties of kombucha mats can further advance ideas on electricity based integration, and possibly, protocognition of symbiotic organisms [36, 37, 38, 39].

Third, kombucha mats, when properly cured, show properties similar to that of clothing materials [40, 41, 42, 43, 44, 45]. In a light of ongoing research on sensing and computing mechanisms embedded in living wearables [46, 47, 35, 48] we aim to evaluate kombucha zoogleal mats as potentially embeddable devices with non-linear and non-trivial electrical properties.

## 2. Methods

The kombucha zooglea was initially obtained from Freshly Fermented Ltd (Lee-on-the-Solent, PO13 9FU, UK) and then produced *in situ*. The infusion was prepared in the following proportions: 2% of tea (PG Tips UK), 5% sugar (Silver Spoon, PE2 9AY, UK), 1 L of boiled tap water. Containers with kombucha were kept at 20-23°C in darkness. The solution was changed every 7 days. An electrical resistance, see scheme and photo of the experimental setup in Fig. 1, of the zoogleal mats was recorded using pairs of platinum round wire 0.50mm diameter (Cookson Precious Metals Ltd., Birmingham, UK) with Flux multimeter 8846A (http://www.fluke.com). Distance between the electrodes was c. 1 cm. We recorded the resistance at one sample per second acquisition frequency. We have conducted over 22 trials of recordings, each one lasted 10^4^ sec (2.77 h), the overall duration of each recording was limited by the factory setting of the Fluke multimeter device. We identified and analysed 90 spikes.

**Figure 1:**
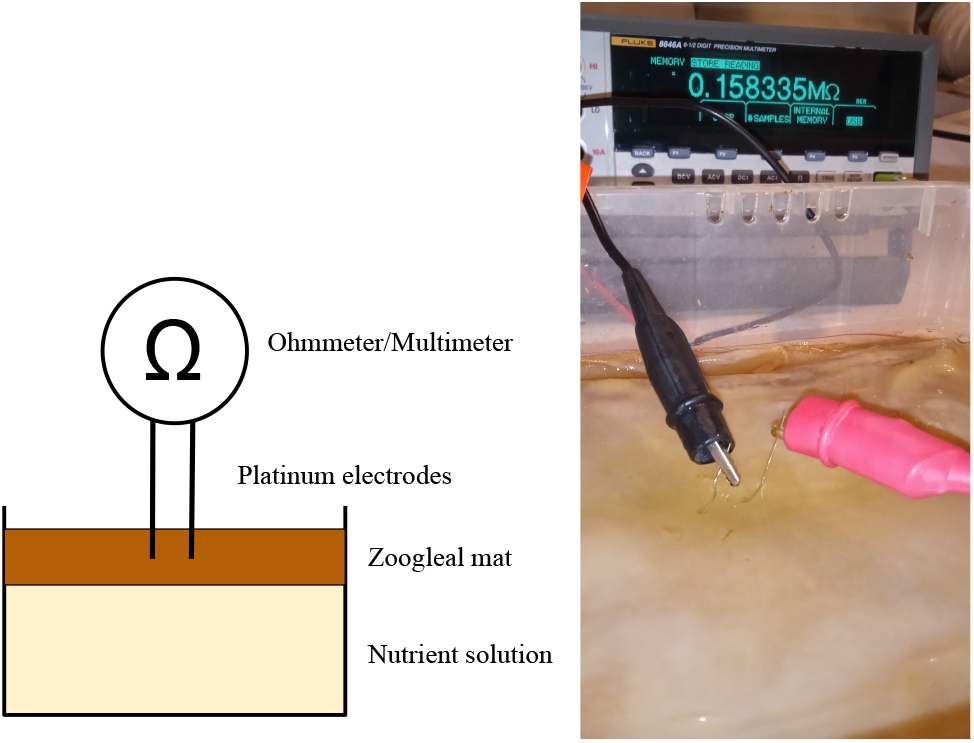
Experimental setup. (a) A scheme. (b) A photo.

## 3. Results

Kombucha zoogleal mats demonstrated rich dynamics of resistance, in terms of morphological complexity, yet the resistance valued remained in a relatively narrow range 0.13 MOhm to 0.19 MOhm (Fig. 2a). In each of 22 recordings we observed spikes of resistance. Typically, there were not more than 5-10 spikes during a period of each recording (Fig. 2b). Many of the spikes show morphological similarity to action potential, indeed, with full understanding that in action potential we observe a variation of a transmembrane potential while in resistance spikes we observe a variation of an electrical resistance, as illustrated in Fig. 2c. In this example, the depolarisation phase occurred in 20 sec, with a resistance change of 16880 Ohm. The repolarisation phase is 23 sec, with a resistance change of 14660 Ohm.

**Figure 2:**
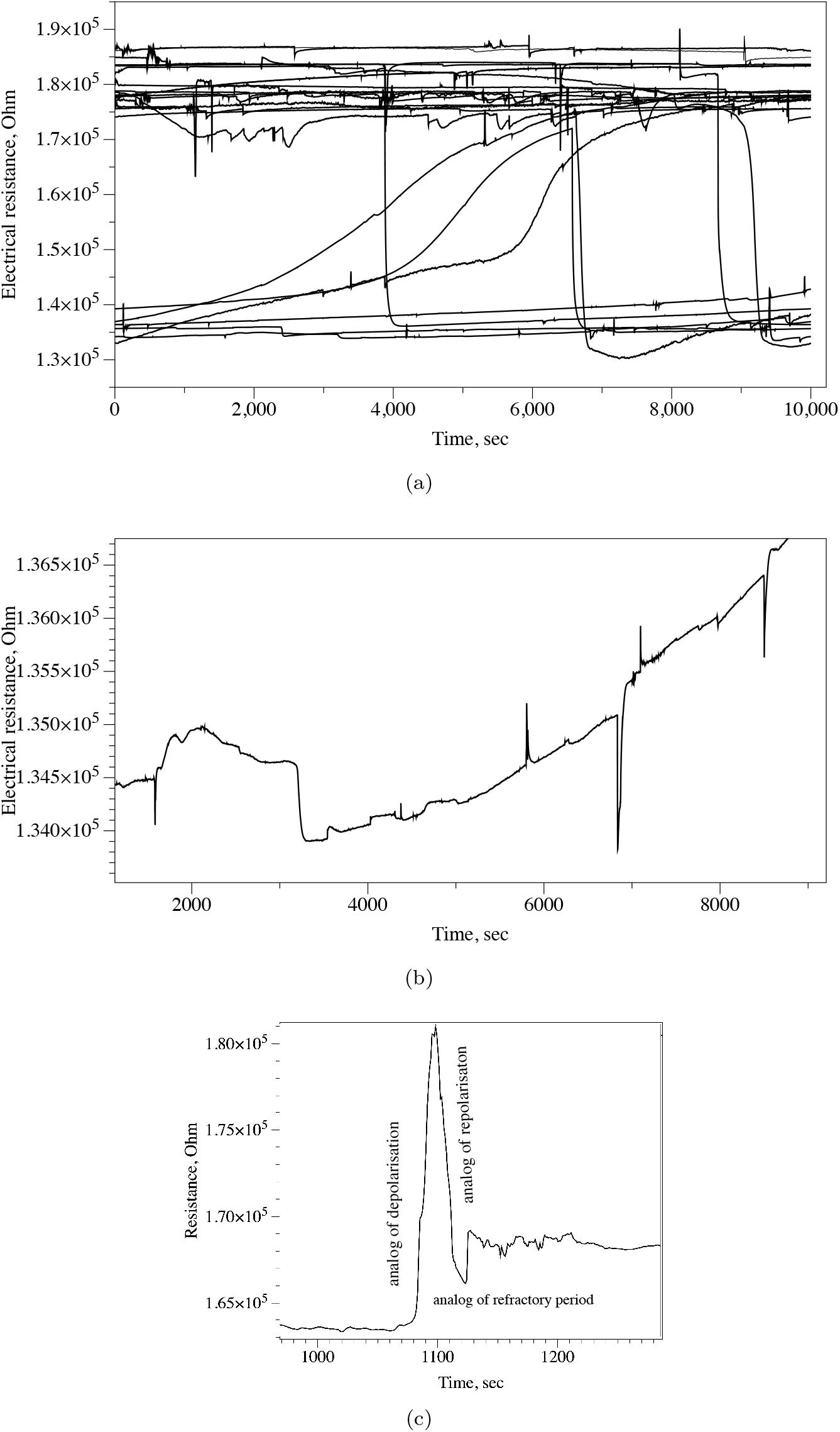
Electrical resistance of kombucha zoogleal mats. (a) All 22 recordings overlapped on one plot to illustrate the diversity of the resistance dynamic. (b) An example of a single recording where solitary spikes are pronounced. (c) Action potential like spike recorded in zoogleal mat.

Spike with versus spike amplitude plot is shown in Fig. 3a. The Pearson correlation coefficient R is 0.0315. Technically speaking the correlation is positive, i.e. wider spikes might have higher amplitude, the relationship between your variables is weak. The coefficient of determination, is 0.001.

**Figure 3:**
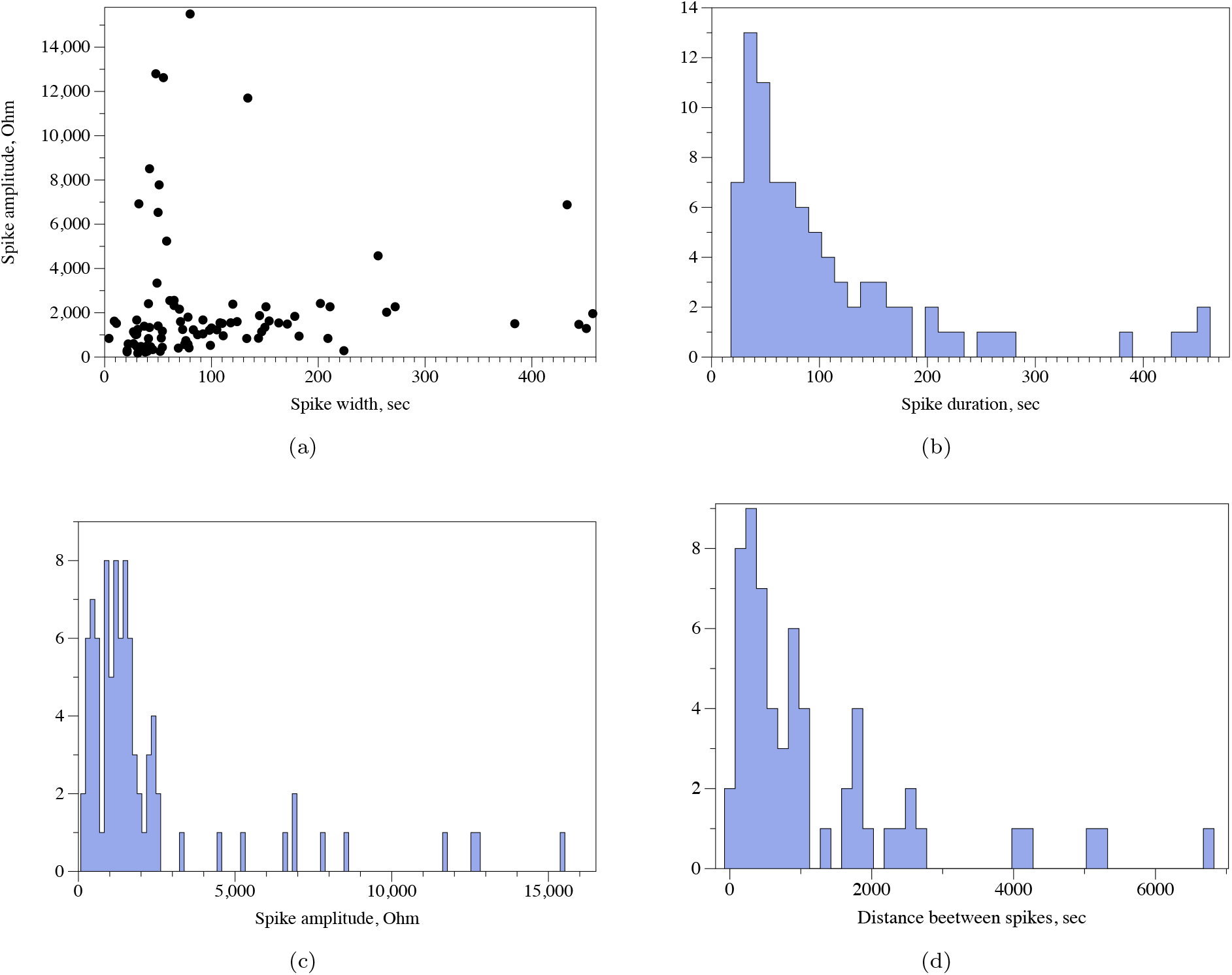
(a) Spike width/duration versus spike amplitude. (b) Frequency distribution of spike width/duration, bin size is 12. (c) Frequency distribution of spike amplitude, bin size is 150. (d) Frequency distribution of distance between spikes, bin size is 150.

Distribution of spike durations is shown in Fig. 3b. The values of the spike duration are in the range 21 sec to 457 sec. Average spike duration is 109.2 sec, standard deviation 99.9, median spike duration is 75 sec. High asymmetry in the distribution is evidenced by skewness 2.03 and kurtosis 4.3.

Frequency distribution of spike amplitudes is shown in Fig. 3c. Minimum amplitude is 172 Ohm and maximum 15500 Ohm. Average amplitude is 2184 Ohm, median 1322 Ohm, standard deviation 2882. Distribution of amplitudes shows higher asymmetry, skewness is 2.8, than distribution of spike duration, longer tail, kurtosis 8.6.

Distribution of values of the distance between spikes is shown in Fig. 3d. An average distance is 1192 sec, median 698 sec, standard deviation 1383, skewness 2.1, kurtosis 4.7.

We define a train of spike is a sequence of spikes where a distance between two neighbouring spikes does not exceed a width/duration of a spike. Twenty two trains of spikes has been identified. An example of a train of five spikes is shown in Fig. 4a. Average width of spikes is 3.6 min (*s* = 1.5), average distance between spikes is 4.8 min (*s* = 1.3), average amplitude is 2173 Ohm (*s* = 289). An example of a solitary (i.e. only this train was observed during 2.77 h recording) train of spikes is shown in Fig. 4. Average width of spikes is 7.2 min (*σ*=1.1 min), average distance between spikes is 10.9 min (*σ*=3.5 min), and average amplitude is 3699.7 Ohm (*σ*=496 Ohm). Length of the train is 28.4 min. Amongst 22 trains, 16 trains had two spikes each, three trains of three spikes each, two trains with four spikes each and one train of five spikes, i.e. majority of trains were comprised of two spikes.

**Figure 4:**
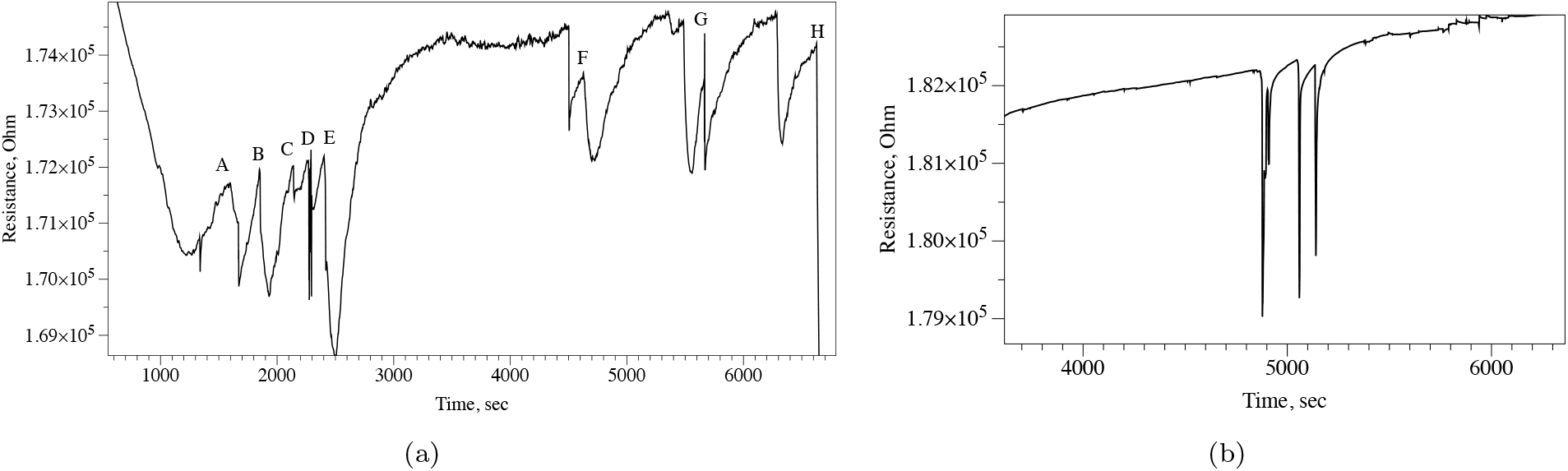
Examples of spiking activity. (a) A train of five spikes A to E, followed by solitary spikes F to H. (b) A solitary train of three spikes.

## 4. Discussion

We found that the electrical resistance of kombucha zooglea mats, symbiotic societies of many species of yeasts and bacteria, exhibits a spiking dynamics. The spikes of electrical resistance are morphologically similar to action potential spikes. Average duration of a resistance spike is 1.8 min, average amplitude is 2.2 kOhm. Average interval between resistance spikes is c. 20 min. In [49] we found that oyster fungi *P. ostreatus* show oscillations of resistance with trains of resistive spiking emerging. Spikes amplitude vary from 1 kOhm to 1.6 kOhm and duration of spikes is 25 min in average. An average distance between spikes is 45 min. Thus, comparing dynamics of electrical resistance of kombucha zoogleal mats and electrical resistance of oyster fungi, we see that the resistive spike amplitudes are of similar ranges. Duration of the oyster fungi spike is nearly 14 times longer than that of kombucha mat while an average distance between resistance spikes in oyster fungi is just a bit over two times longer than that in kombucha mats.

What is an underlying mechanism of the resistance spiking?

Whilst we could not provide any firm propositions, our vision is that the resistance spikes might be byproducts of the electrical potential spiking in kombucha mat. Let us compare parameters of kombucha mats’ electrical potential spiking analysed in [33] with parameters of electrical resistance spiking. There are 3.2 electrical potential spikes in average in a train of spikes, median 3. There are 2.4 electrical resistance spikes, median 2. In fastest trains of electrical potential spikes, a spike width is 2.8 min, median 3.2 min while average duration of an electrical resistance spike is 1.8 min, median 1.2 min. Thus the parameters of electrical potential spikes and electrical resistance spikes differ just by c. 30% and are of the same range.

This all indicates that, possibly, travelling waves of electrical activity in kombucha mats change biochemical composition of the mats to temporarily modify their resistance. As we discussed in [33], electrical potential spiking in kombucha mats might be due the waves of depolarisation, emerging due to metabolically triggered release of potassium, travelling in zoogleal mats. An evidence of such travelling waves, albeit in bacterial films, is provided in [50].

Another possible reason for the resistance spikes could be related to glycolytic oscillations of yeasts, which are reflected in oscillations of their resistance and capacitance [51]. However, periods of glycolytic oscillations are of an order of hours, not minutes as in kombucha mats resistance spikes.

One more explanation of the resistance spikes might in the fact that living cells and their colonies can be treated as leaking condensers [52, 1]. These condensers can discharge in an oscillator manner [53], their discharges might affect local flow of ions, thus leading to spikes in the electrical resistance. What would be further directions of research into electrical resistance dynamics of kombucha zoogleal mats?

First, we can attempt to develop bio-hybrid sensors. Previously, we demonstrated that a dynamics of electrical potential of kombucha is changing after optical, chemical and electrical stimulation [33]. If a dynamic of electrical resistance is somehow linked to the dynamic of electrical potential, we could propose that a stimulation of kombucha zoogleal mat might lead to changes in their resistance. Then it will be feasible to integrate kombucha with bio-sensing devices, because detecting changes in resistance is much simpler than detecting changes in electrical potential spiking.

Second, non-trivial dynamics of electrical resistance of kombucha mats indicates possible memristive [54, 55], or at least memfractive [56, 57], properties. Therefore, it would be possible to implement neuromorphic architectures and memristive computing devices [58] with living kombucha mats, thus opening a new era in soft self-repairable bio-electronic devices. A very low frequency of kombucha zooglea electronic oscillators does not preclude us from considering inclusion of the electrical resistance oscillators in bio-hybrid analog circuits embedded in living wearables [47, 35, 59, 60].

## 5. Acknowledgement

This project has received funding from the European Union’s Horizon 2020 research and innovation programme FET OPEN “Challenging current thinking” under grant agreement No 101071145.

## 6. Availability of Data and Materials

The datasets used and/or analysed during the current study available from the corresponding author on reasonable request.

## Notes

### Competing Interest Statement

The authors have declared no competing interest.

